# The SMC complex, MukBEF, organizes the *Escherichia coli* chromosome by forming an axial core

**DOI:** 10.1101/696872

**Authors:** Jarno Mäkelä, David J. Sherratt

## Abstract

Structural Maintenance of Chromosomes (SMC) complexes organize and individualize chromosomes ubiquitously, thereby contributing to their faithful segregation. Here we explore how *Escherichia coli* chromosome organization emerges from the action of the SMC complex MukBEF, using quantitative imaging in cells with increased MukBEF occupancy on the chromosome. We demonstrate that the *E. coli* chromosome is organized as series of loops around a thin axial MukBEF core whose length is ~1100 times shorter than the chromosomal DNA. The core is linear (1 μm), or circular (1.5 μm) in the absence of MatP, which displaces MukBEF from the 800 kbp replication termination region (*ter*). Our findings illustrate how MukBEF compacts the chromosome lengthwise and demonstrate how displacement of MukBEF from *ter* promotes MukBEF enrichment with the replication origin.

## Main Text

MukBEF, the *Escherichia coli* SMC complex homolog, exhibits the distinctive SMC complex architecture, with the additional feature that the kleisin, MukF, is dimeric, leading to the formation of dimer of dimer complexes *in vivo* and *in vitro* (Fig. 1A) (*1*–*4*). Impairment of MukF dimerization leads to a failure of MukBEF complexes to stably associate with the chromosome (*5*). MukBEF homologs, containing a dimeric kleisin, are confined to γ-proteobacteria and have co-evolved with a set of genes, including MatP, which binds to ~23 *matS* sites in the ~800 kb chromosome replication terminus region (*ter)* (Fig. 1B)(*6*, *7*). Interaction of MukBEF with MatP-*matS* leads to displacement of MukBEF complexes from *ter* (*8*). Furthermore, MukB interacts with the chromosome decatenase, topoisomerase IV, providing a functional link between chromosome organization and unlinking (*8*–*10*). MukBEF impairment leads to defects in chromosome segregation and anucleate cell production (*11*, *12*). Clusters of positioned MukBEF complexes form ‘foci’ by fluorescence microscopy in wild-type cells (Fig. 1C)(*2*, *8*, *12*). These clusters position replication origin regions (*oriC*) to either mid-nucleoid (new-born cells) or the nucleoid quarter positions (cells that have replicated and segregated their *oriC* regions), with the MukBEF clusters being positioned on the nucleoid by a Turing patterning mechanism (*2*, *13*, *14*). ~50% of ~200 MukBEF complexes/cell are tightly associated with DNA (residence time ~50 s), while 25-50% of these are present in foci (*2*).

**Fig. 1.**
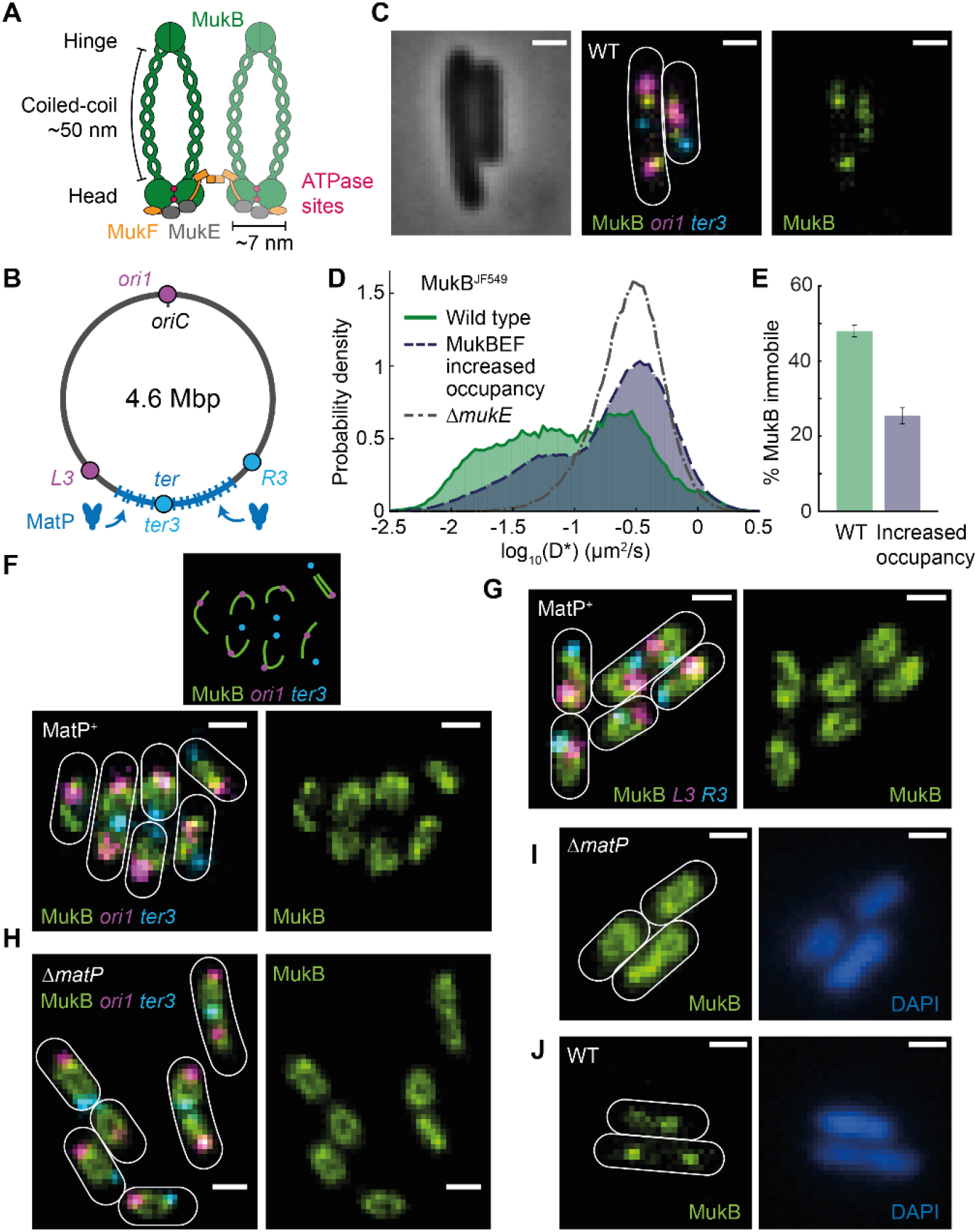
MukBEF architecture and function. (**A**) MukBEF architecture. (**B**) *E. coli* chromosome showing 800 kbp *ter* region with *matS* sites (blue bars) that are specifically bound by MatP. Locations of *ori1*, *ter3*, *L3*, and *R3* markers are shown. (**C**) Representative phase contrast and fluorescence images of wild type cells with labeled MukB, *ori1*, and *ter3*. Scale bar, 1 μm. (**D**) Log-scale distribution of apparent diffusion coefficients (D*) from single molecule tracking of MukB^JF549^ in wild type (38502 tracks), MukBEF increased occupancy (355560 tracks) and Δ*mukE* (166991 tracks) cells. Data are from 3 repeats. (**E**) Percentage of immobile MukB tracks (D* < 0.0875 μm^2^/s) in wild type and MukBEF increased occupancy cells. The threshold was defined by the lower 5% quantile of D* in Δ*mukE* cells. Error bars denote SEM. (**F**)-(**J**) Representative fluorescence images with cell borders shown (**F**) MukBEF increased occupancy cells with *ori1* and *ter3*, (schematic on top) (**G**) MukBEF increased occupancy cells with *L3* and *R3*, (**H**) MukBEF increased occupancy Δ*matP* with *ori1* and *ter3*, (**I**) MukBEF increased occupancy Δ*matP* cells, and (**J**) wild type cells with DAPI-stained nucleoids. Scale bars, 1 μm.

To address how MukBEF organizes the *E. coli* chromosome, and why MukBEF clusters colocalize with *oriC*, we undertook a quantitative and super-resolution imaging analysis of *E. coli* cells which had an increased MukBEF occupancy on the chromosome as a consequence of 6.3 ± 0.4-fold over-expression from the endogenous *mukBEF* operon, after replacing the native promoter with an inducible promoter, *P*_*ara*_ (Fig. S1). The occupancy of bound MukBEF complexes in wild type and over-expressing cells was determined by single molecule tracking using a functional HaloTag fusion to the endogenous *mukB* gene (Fig. 1D)(*15*). MukBEF over-expression cells had 25.4 ± 2.1% of molecules immobile (chromosome-associated) as compared to 48 ± 1.5% molecules in wild type cells (Fig. 1E). Therefore, the increase in occupancy on the chromosome was ~3-fold. Increased MukBEF occupancy cells had the same generation time and percentage of anucleate cells as wild type cells (Fig. S1). Additionally, we determined almost identical residency times for the chromosome-associated MukBEF complexes in normal (64 ± 14 s) and increased occupancy cells (67 ± 15 s)(Fig. S1), similar to the estimate for wild type complexes using FRAP (*2*).

Under conditions of increased occupancy, fluorescent MukBEF formed a filamentous axial core that was predominantly linear (cells with MatP present, MatP^+^) or circular (Δ*matP* cells)(Fig. 1F, G, H, and Fig. S2). A *ter3* marker, located close to the center of *ter*, was depleted of MukBEF fluorescence in wild type, but not in Δ*matP* cells, demonstrating that the difference in axial core structure is a direct consequence of the MatP-*matS*-dependent displacement of MukBEF from *ter*. MukBEF does not link chromosome arms together, instead, the ends of linear axial cores in MatP^+^ cells localized near *L3* and *R3* markers that flank the *ter* region (Fig. 1A, G). DAPI-stained nucleoids of increased MukBEF occupancy cells showed no detectable morphological differences to those in wild type cells (Fig. 1H, I, Fig. S3), consistent with overall chromosome compaction unaffected by increased occupancy. MukBEF axial cores were also observed in rifampicin-treated cells (reduced molecular crowding) and in cells of increased volume after treatment with A22 (Fig. S4)(*16*).

To quantitatively analyze MukBEF axial cores in relation to genetic markers, we enriched for cells with completely replicated chromosomes by incubation with serine hydroxamate (SHX), thereby avoiding bias from partially replicated chromosomes. In single chromosome MatP^+^ cells, the brightest MukBEF pixel to *ori1* distance (increased occupancy 0.24 ± 0.01 μm; WT 0.26 ± 0.002 μm) was much smaller than to *ter3* (increased occupancy 0.52 ± 0.02 μm; WT 0.59 ± 0.004 μm) (Fig. 2A, B). In single chromosome Δ*matP* cells, the brightest MukBEF pixel distances were nearly identical between *ori1* (increased occupancy 0.34 ± 0.003 μm; WT 0.29 ± 0.01 μm) and *ter3* (increased occupancy 0.37 ± 0.001 μm; WT 0.34 ± 0.02 μm) (Fig. 2C, D). Therefore, even though the MukBEF foci in wild type cells are replaced by the axial cores after increasing MukBEF occupancy, enrichment of MukBEF with *ori* remains. In contrast, circular axial cores in increased occupancy Δ*matP* cells were uniform with respect to *ori1*/*ter3* loci (Fig. 2C). Next, to assess how increased chromosome occupancy in replicating cells reflects the behavior of all chromosome-associated MukBEF, we determined the normalized intensity of every cellular MukBEF pixel as a function of distance to *ori1* or *ter3* (Fig. 2E, F). In MatP^+^ cells, the MukBEF intensity accumulates in the vicinity of *ori1*, far from *ter3*, while Δ*matP* cells exhibited similar descending profiles for both *ori1* and *ter3*. Importantly, the intensity profile patterns were almost identical in increased occupancy cells, in wild type cells, and in cells during the induction period of MukBEF over-expression (Fig. S5). These observations indicate that the nature of association of MukBEF with chromosomes is unaffected by increased MukBEF occupancy; while less of the chromosome is occupied by MukBEF complexes at any given time in wild type cells, the probability of MukBEF occupying a chromosome locus remains the same.

**Fig. 2.**
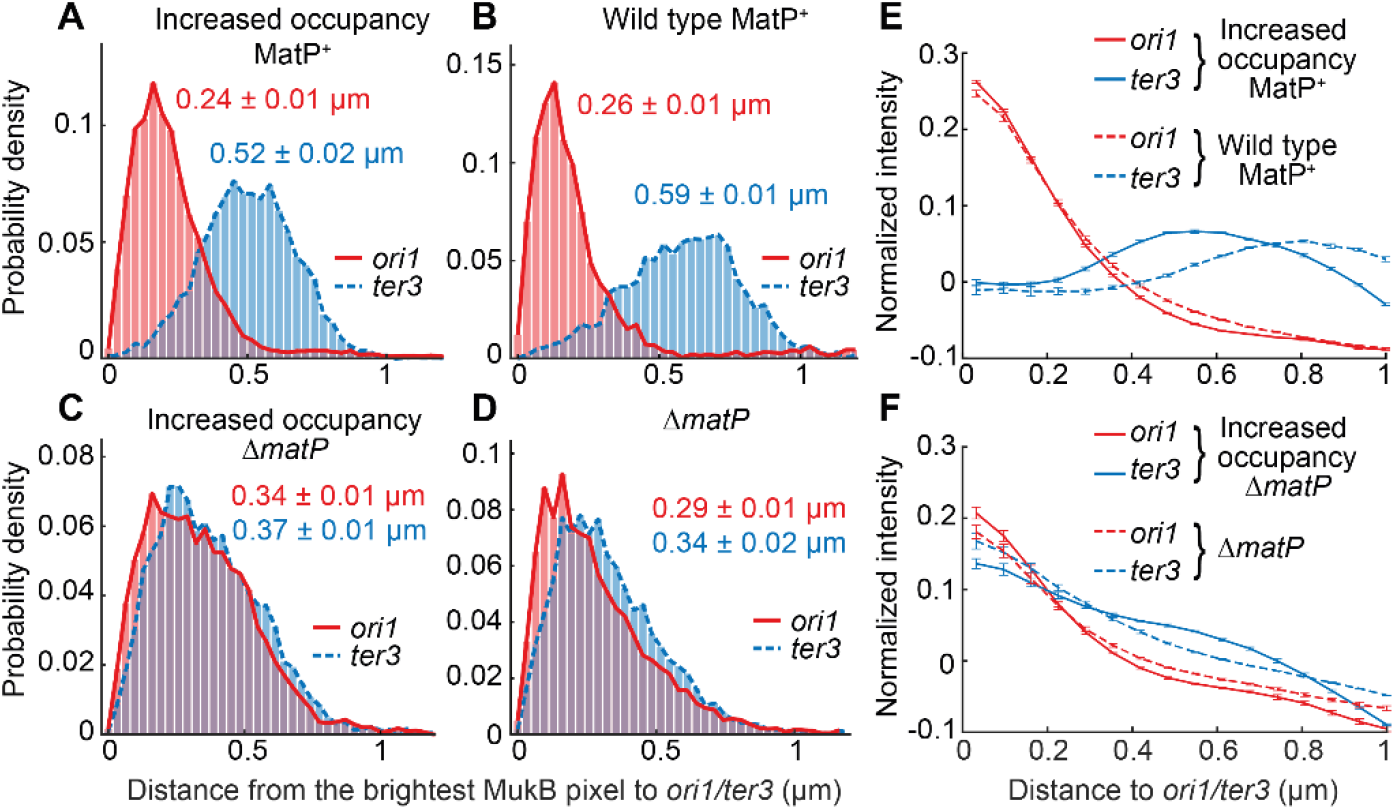
Quantitative analyses of MukBEF localization. (**A**)-(**D**) Distances between the brightest MukB pixel and *ori1*/*ter3* markers (±SEM) in (**A**) MukBEF increased occupancy, (**B**) wild type, (**C**) MukBEF increased occupancy Δ*matP* and (**D**) Δ*matP* cells. Prior to imaging cells were treated with SHX and only cells with a single chromosome were included in the analysis. Data are from 3 repeats. (**E**)-(**F**) Normalized MukB pixel intensity as a function of distance to *ori1*/*ter3* in asynchronous populations of (**E**) MatP^+^ and (**F**) Δ*matP* cells with wild type and increased MukBEF occupancy. Error bars, SEM from 3 repeats.

To gain more insights into the nature of the axial core in live cells, three-dimensional structured illumination microscopy (3D-SIM) was used (Fig. 3A, B, C, and Fig. S6). The axial cores were better resolved than those by epifluorescence microscopy, but had the same overall appearance (compare Fig. 1F, G, H). To estimate the lengthwise compaction of the chromosome by MukBEF, the length of the axial cores was quantified in cells with completely replicated chromosomes. Their contour length in increased occupancy Δ*matP* cells was 1.45 ± 0.01 μm (Fig. 3D). Given that the 4.64 Mbp circular chromosome of *E. coli* has a contour length of 1.58 mm, this corresponds to a ~1100-fold lengthwise compaction. In increased occupancy MatP^+^ cells, the linear axial cores were 1.03 ± 0.06 μm long (Fig. 3E), extending from genetic markers *L3* to *R3*, which bound the *ter* region (3.22 Mbp; 69.5% of the genome). Since *L3* and *R3* colocalize with the ends of the axial core, the displacement of MukBEF by MatP-*matS* extends beyond the outer *matS* sites. Consequently, the overall level of compaction (~1100-fold) in the linear axial cores is similar to that of the circular cores in Δ*matP* cells. Finally, the peak of measured thickness distribution of the axial cores was ~130 nm (Fig. 3F), close to the limit of SIM resolution (~120 nm); thereby suggesting that the actual thickness is likely less and of the same order as the dimension of functional MukBEF complexes (Fig. 1A). In combination with the estimated MukBEF increased occupancy numbers on the chromosome (Fig. 1E, Fig. S1)(*2*), we infer that there is one dimer of dimers complex for every ~6 nm of axial core length, consistent with it being a near continuous array of MukBEF complexes. We also estimate the length of DNA per MukBEF dimer of dimers to be ~22 kbp. To gain insights into the overall compaction of MukBEF displaced region that includes 800 kbp *ter* region, we measured the minimal length between *L3* and *R3* markers (1.42 Mbp) in MatP^+^ (Fig. 1B). We observed a bimodal length distribution, indicating that this region can be either more loosely (~400-fold), or densely, (~1000-fold) compacted (Fig. S7). This is consistent with the observation that different genetic markers in *ter* can localize to distant regions of the same cell (*17*), with Hi-C data (*18*), and with analysis of cells with increased volume (*16*). We summarize the dimensions of MukBEF axial filaments in Fig. 3G.

**Fig. 3.**
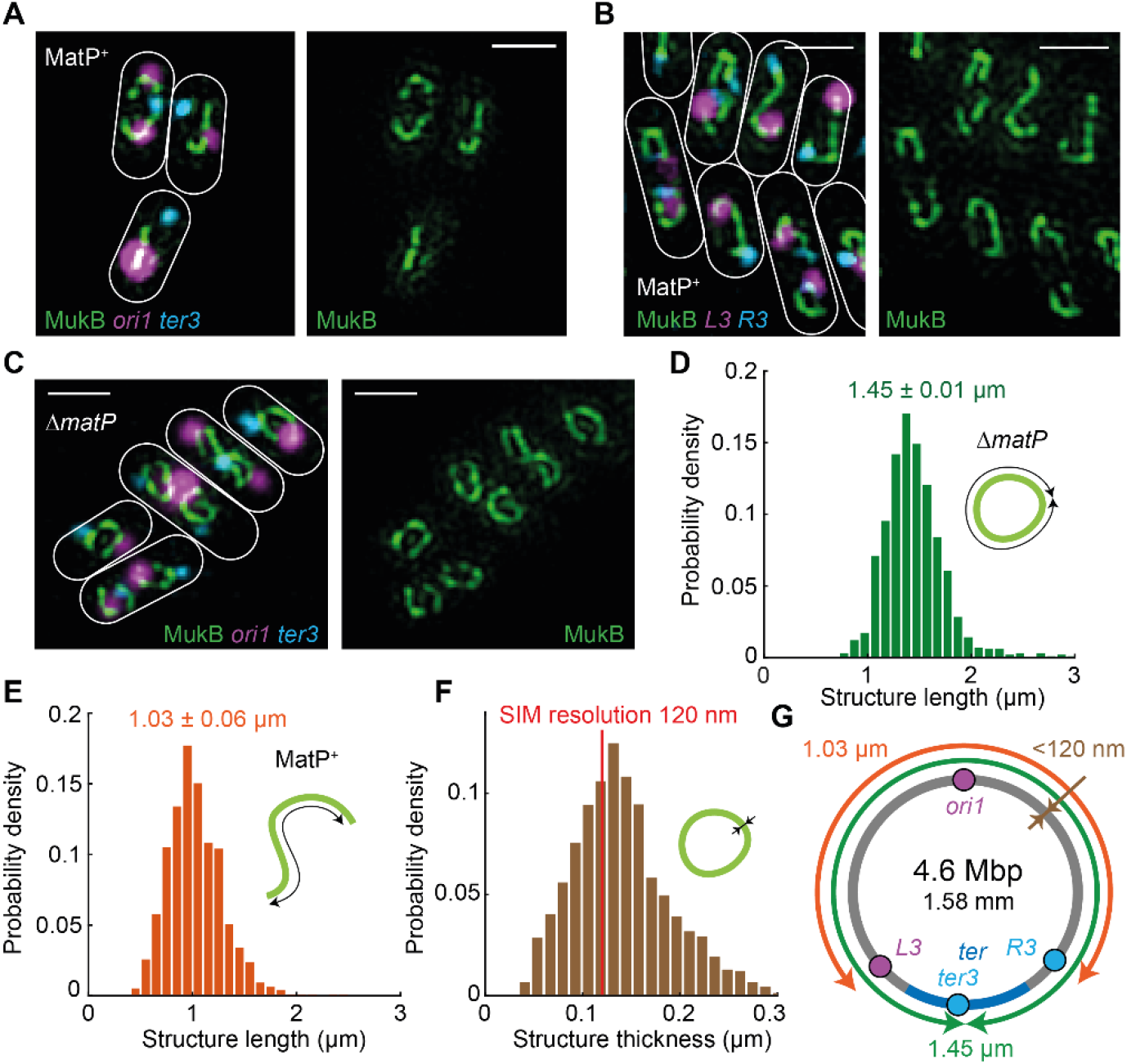
3D-SIM analysis of MukBEF axial cores. (**A**)-(**C**) Representative SIM images of MukBEF increased occupancy in (**A**) MatP^+^ with *ori1* and *ter3*, (**B**) MatP^+^ with *L3* and *R3* and (**C**) Δ*matP* with *ori1*, *ter3* cells. 3D-SIM images were projected onto 2D for visualization. Scale bars, 1 μm. (**D**) Distribution of axial core contour lengths (*n* = 986, ±SEM) measured from MukBEF increased occupancy Δ*matP* cells. (**E**) Distribution of linear axial core lengths (*n* = 971, ±SEM) measured from MukBEF increased occupancy cells. (**F**) Distribution of MukBEF axial core thickness (*n* = 6657, ±SEM). The peak is at 132 nm. Red line denotes the resolution of SIM. Prior to imaging cells were treated with SHX to prevent replication initiation. Data are from 3 repeats. (**G**) Schematic of MukBEF axial core dimensions.

MukBEF simply capturing two random segments of DNA cannot generate the observed axial cores (Fig. 4A). As a model, progressive enlargement of loops promotes linear chromosome compaction and formation of brush-like chromosome (*19*). To understand how loop extrusion can lead to the formation of the observed MukBEF axial cores, we undertook stochastic simulations of the process using experimentally derived MukBEF parameters (Fig. 4B). We modeled MukBEF complexes as dimers of dimers that load randomly onto the chromosome, and are capable of extruding loops bidirectionally at a rate of 600 bp/dimer/s (*20*). We assume dimers of dimers cannot overtake each other, and that when loop extrusion brings two translocating complexes together, the inner dimers stall, while the outer dimers continue to extrude loops. MukBEF complexes spontaneously dissociate from chromosomes after an exponential dwell time (65 s), while any complexes that encounter MatP-*matS* in *ter* are immediately released from DNA. Representative simulations are shown with (MatP^+^) and without (Δ*matP*) MukBEF displacement from *ter* (Fig. 4C). As MukBEF occupancy on the chromosome increases, more of the chromosome is included into MukBEF-generated loops (wild type 48%; increased occupancy 72%) (Fig. 4D), while the loop size generated by individual MukBEF complexes decreases (wild type 52 kbp; increased occupancy 31 kbp), due to more frequent collisions during loop extrusion (Fig. 4E). Nevertheless, colliding MukBEF complexes form continuous clusters that can contain up to 800 kbp of DNA with wild type occupancy or >1 Mbp with increased occupancy (Fig. 4F, Fig. S8). By comparison, unidirectional loop extrusion was found inefficient (Fig. 4D, F, and Fig. S8), as inferred elsewhere (*21*). The requirement for MukBEF to act as dimers of dimers, provides a plausible mechanism for bidirectional loop extrusion, while the *in vivo* significance of unidirectional loop extrusion by condensin in single-molecule *in vitro* experiments (*20*) remains unclear. Experimentally, the MukBEF axial core is more lengthwise compacted (~1100-fold) than the simulations predict (~10-fold). Therefore, we propose that while MukBEF is responsible for forming the chromosome axial core from which loops of different size emanate, other looping proteins and processes contribute further to the overall lengthwise compaction. We expect wild type MukBEF axial cores to be more granular, less continuous entities, but with a comparable level of chromosome compaction.

**Fig. 4.**
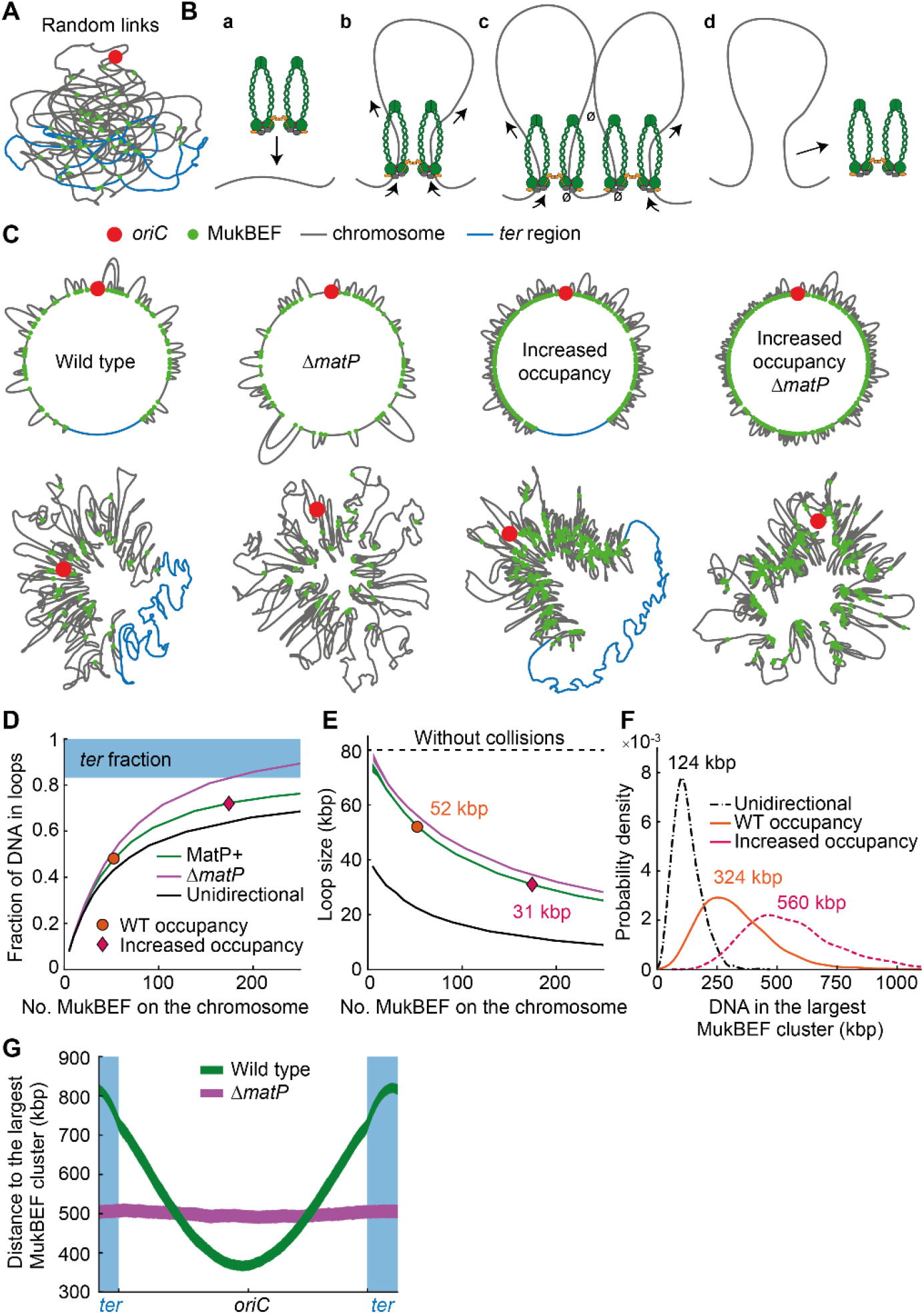
Modeling loop extrusion by MukBEF. (**A**) Example of a randomly linked *E. coli* chromosome. (**B**) Description of the model: (**a**) A MukBEF dimer of dimers associates at a random site on the chromosome. (**b**) The loop is enlarged by the two dimers moving in opposite directions along the chromosome. (**c**) Collisions between MukBEFs prevent loop extrusion only on the internal collided dimers. (**d**) MukBEF spontaneously dissociates releasing the loop. *ter* region immediately displaces MukBEF upon contact. (**C**) Representative *E. coli* chromosomes with/without MukBEF displacement from *ter* and wild type/increased MukBEF occupancy. (top) Beginning and end of loops with MukBEF (green dots) along the chromosome. (bottom) Force-directed layouts of the chromosomes (see SI). (**D**) Fraction of the chromosome within a loop and (**E**) loop size per individual MukBEF as function of the number of chromosome-bound MukBEF dimers of dimers. In unidirectional loop extrusion, dimers are not connected and each dimer acts independently, binding and extruding a loop at randomly chosen direction. Experimentally estimated numbers of bound MukBEF dimers of dimers for wild type and increased occupancy cells are shown. (**F**) Distribution of DNA in the largest MukBEF cluster (no unlooped DNA between them) for wild type and increased MukBEF occupancy, and unidirectional loop extrusion (see also Fig. S8). (**G**) Shortest distance from a chromosome locus to the largest MukBEF cluster along the chromosome with wild type occupancy and with/without MukBEF displacement from *ter*. Line thickness denotes 95% bootstrap confidence interval for the mean across 1000 simulation replicas.

We finally considered whether the model explains the enrichment of MukBEF at *ori1* (Fig. 2B). We computed the distance along the chromosome with wild type occupancy from a locus to the largest MukBEF cluster and, similar to experiments, we observed a minimum distance between the largest MukBEF cluster and the *oriC* region (Fig. 4G). This is a consequence of MatP depleting MukBEF from *ter* (Fig. S8) that is opposite to the *oriC* in the center of the MukBEF occupied region. The absence of MukBEF displacement from *ter* abrogates this effect. In agreement, we showed that MukBEF clusters colocalized almost equally with *ori1* and *ter3* in Δ*matP* cells (Fig. 2D). We note that this is not dependent on loop extrusion *per se* and other linear chromosome compaction mechanisms can similarly reproduce the colocalization. Two contrasting strategies for preferential association of SMC complexes with bacterial *oriC* regions have been identified. Those bacteria that have MukBEF along with MatP-*matS*, use displacement of MukBEF from *ter*, while most characterized bacteria preferentially load their SMC complexes directly to *oriC*-proximal ParB-*parS* sites (*22*, *23*).

The MukBEF axial cores are at least superficially similar to ‘vermicelli’ chromosomes observed in mammalian cells when cohesin occupancy was increased by impairing WAPL, which stimulates ATP hydrolysis-dependent cohesin removal from chromosomes (*24*). Furthermore, the MukBEF axial cores resemble the axial scaffolds formed by condensin II during the early stages of mitotic chromosome formation (*25*). Individualization and separation of chromosome arms by MukBEF contrasts to the situation in *Bacillus subtilis*, where the action of the SMC complex zips-up the two chromosome arms, rather than separating them (*23*), although we surmise whether the putative SMC-directed loops could form within a chromosome arm, with higher order interactions bringing the two arms together. Nevertheless, in *B. subtilis*, and other studied bacteria that have canonical SMC complexes, the two chromosome arms are colinear along the cell long axis, contrasting with the situation in *E. coli*, and perhaps other MukBEF-MatP-containing bacteria (*26*). The MukBEF interaction with the decatenase, TopoIV, means that the patterns of MukBEF association with chromosomes is mimicked by those of TopoIV, with its action being enriched close to *oriC* and depleted from *ter*, with coupling between the activities of both enzymes possible (*8*). We conclude that chromosome-associated MukBEF complexes are the template and ‘machine’ for formation of DNA loops in chromosomes, and their characterization adds weight to the hypothesis that lengthwise compaction by intra-chromosome loop formation is the mechanism by which all SMC complexes organize and individualize chromosomes.

## Acknowledgments

The Micron imaging unit provided microscopes and Lothar Schermelleh provided advice and expertise with SIM. We thank other members of the Sherratt and Uphoff groups, Katarzyna Ginda-Mäkelä, Sophie Nolivos, Bela Novak and Kim Nasmyth for insightful discussions. Sophie Nolivos undertook early experiments on the interplay between MukBEF-MatP-*matS*. Luke Lavis (Janelia Farm) provided the Halo ligand dye JF549. This work was funded by Wellcome (Investigator Award to DJS; 200782/Z/16/Z).

## Author contributions

The research was devised and directed by JM and DJS. JM undertook and analyzed experiments, and did the simulations. JM and DJS wrote the paper.

## Materials and Methods

### Bacterial strains and growth

#### Strain construction

Bacterial strains and primers are listed in Tables S1 and S2, respectively. All strains were derivatives of *E. coli* K12 AB1157 (*27*). To replace the native *smtA-mukBEF* promoter with an inducible *araC-P*_*BAD*_, first, a plasmid containing *kan-araC-P*_*BAD*_ in the pBAD24 (*28*) was constructed. Subsequently, *kan-araC-P*_*BAD*_ was PCR amplified and the product used to replace the native promoter of *smtA-mukBEF* operon in strain SN192 with λ-red recombination (*29*). Finally, the generated chromosomal gene locus was transferred by phage P1 transduction to SN192 yielding strain JM90. P1 transduction was also used to introduce *kan-araC-P*_*BAD*_*-smtA-mukFE-mukB-Ypet* into SN302, and SN191 resulting in strains JM91, and JM101, respectively. JM103 was constructed by first removing the *kan* resistance gene from JM90 using Flp recombinase from plasmid pCP20 (*29*), introducing *mukB-haloTag-kan* from JM41 by λ-red recombination and transferring the generated chromosomal gene loci into AB1157 by P1 transduction. The JM56 strain (*mukB-haloTag*, Δ*mukE*) was constructed by removing the *kan* resistance gene from JM41 using Flp recombinase and replacing the endogenous *mukE* gene with a kanamycin cassette using λ-red recombination. All genetic modifications were verified by PCR, the Muk^+/−^ phenotype was verified by temperature-resistance or lack of it in rich media, and behavior in quantitative imaging, as described (*8*).

#### Growth conditions

Cells were grown in M9 minimal medium supplemented with 0.2% (v/v) glycerol and required amino acids (threonine, leucine, proline, histidine and arginine - 0.1 mg ml^−1^) at 30 °C. For MukBEF over-expression strains, cells were additionally grown with a constant presence of 0.2% (w/v) L-(+)-Arabinose. For microscopy, cells were grown overnight, diluted 1000-fold and grown to an OD_600_ of 0.05–0.2. Cells were then pelleted, spotted onto an M9 glycerol 1% (w/v) agarose pad on a slide and covered by a coverslip. For PALM microscopy, 0.17 mm thickness coverslips were plasma-cleaned of any background fluorescent particles before use.

For experiments in which cells were enriched for completed non-replicating chromosomes, cells were treated with DL serine hydroxamate (SHX) (Sigma-Aldrich, S4503, final concentration of 1 mg ml^−1^). During the treatment, cells do not initiate new rounds of replication, but most complete any ongoing rounds (*30*). To allow sufficient time for ongoing replications to complete, cultures were grown in the presence of SHX for 3 h prior to imaging. This facilitated analysis of MukBEF axial cores and their distance relationships to genetic markers, because ongoing replication can bias results towards smaller *ori1* distances, as the number of *ori1* is greater than number of *ter3* (MatP^+^, *ori1*/*ter3* ratio 1.6; Δ*matP*, *ori1*/*ter3* ratio 1.5).

### Live-cell microscopy experiments

#### Epifluorescence microscopy

Fluorescence images were acquired on an inverted fluorescence microscope (Ti-E, Nikon) equipped with a perfect focus system, a 100× NA 1.4 oil immersion objective, a motorized stage, an sCMOS camera (Orca Flash 4, Hamamatsu), and a temperature chamber (Okolabs). Exposure times were 150 ms for mCherry, and 100 ms for mYpet using an LED excitation source (Lumencor SpectraX).

Cell outlines, fluorescence intensities and FROS marker locations were detected using SuperSegger (*31*). For fluorescence intensity profiles as a function of *ori1*/*ter3* (Fig. 2E, F, Fig. S4), cell pixel intensities were normalized by subtracting the average cell intensity and dividing by the maximum intensity of a cell. Distances of each pixel to the closest *ori1* and *ter3* markers were measured and the average intensity for each binned distance was calculated.

For anucleate cell percentages and DAPI intensity profile analysis, cells were incubated with 1 μg/ml DAPI. For DAPI profiles (Fig. S3), fluorescence intensity along the long cell axis for each cell was extracted. Only cells below 2.6 μm long were considered to avoid cells with more than one chromosomes. Both cell length and intensity were normalized, and these profiles were overlaid. DAPI area length was measured as full-width half-maximum of the DAPI profile. All data analysis was performed in MATLAB (MathWorks).

#### Photoactivated localization microscopy

Live cell single-molecule-tracking photoactivated localization microscopy (PALM) was performed on a custom-built total internal reflection fluorescence (TIRF) microscope built around the Rapid Automated Modular Microscope (RAMM) System (ASI Imaging) with a motorized piezo stage, a z-motor objective mount, and autofocus system (MS-2000, PZ-2000FT, CRISP, ASI Imaging). MukB-HaloTag labelled with JF549 HaloTag ligand was measured with a 100 mW 561 nm laser with 15% transmission (iChrome MLE, Toptica). The laser was collimated and focused through 100× oil immersion objective (NA 1.4, Olympus) onto the sample using an angle for highly inclined thin illumination (*32*). Fluorescence emission was filtered by a dichroic mirror and notch filter (ZT405/488/561rpc and ZET405/488/561NF, Chroma). Fluorescence emission was measured using an EMCCD camera (iXon Ultra, 512×512 pixels, Andor) with a pixel size of 96 nm. Transmission illumination was provided by an LED source and condenser (ASI Imaging). PALM movies were acquired with a frame time of 15.48 ms (*15*).

Single molecule tracking data was analyzed using a custom-written MATLAB software (MathWorks) as in (*15*, *33*). Cell outlines were detected as in the previous section. Fluorescently-labelled MukB were detected by using band-pass filtering and an intensity threshold to each frame of the movie. These initial localizations positions were used as a start point in a two-dimensional elliptical Gaussian fit for high-precision localization. Fitting parameters were x-position, y-position, x-width, y-width, elliptical rotation angle, intensity, and background. Single molecule tracking was performed by linking positions to a track if they appeared in consecutive frames within a window of 0.48 μm as in (*33*). In rare cases of multiple localizations within the tracking radius, tracks were linked such that the sum of step distances was minimized. Tracking allowed for a transient (1 frame) disappearance of the molecule within a track due to blinking or missed localization. The mobility of each molecule was determined by calculating an apparent diffusion coefficient, D*, from the stepwise mean-squared displacement (MSD) of the track using (*33*):

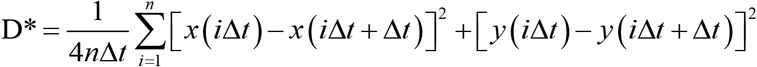

where *x(t)* and *y(t)* are the coordinates of the molecule at time *t*, the frame time of the camera is Δ*t*, and *n* is the number of the steps in the trajectory. Tracks shorter than *n* = 4 steps long were omitted due to the higher uncertainty in D*. Threshold between mobile and immobile tracks were selected by measuring D* in Δ*mukE* strain (JM56) that does not stably associate with the chromosome and setting threshold to the lower 0.05 quantile of the D* distribution. Below this, threshold molecules were considered to be associated with the chromosome.

#### Measuring long-lasting binding events

PALM movies to measure long duration binding events of MukB-HaloTag labelled with JF549 dye were recorded using 1 s exposure times and low continuous 561 nm excitation (0.1% transmission) that blurs mobile molecules into the background whereas immobile molecules still appear as a diffraction-limited spot. Single molecule localization and tracking was used as described in the previous section. Additionally, bound and mobile molecules were distinguished by the width of the elliptical fits, with a short axis-width < 160 nm and long axis-width < 200 nm to determine bound molecules (*33*) and missing frames were not allowed. The lengths of immobile tracks were measured and a survival probability curve (1-CDF) was shown in Fig. S1D. To extract exponential-time constants, the survival probability curve of the immobile molecules was fitted to a double-exponential function corresponding to specific and non-specific DNA binding (*34*, *35*). A single-exponential function was found to not properly fit the survival probability curve. The fitting was performed using least squares criterion with a weight 1/y to compensate for small values in the tail. The duration of specific DNA binding events was defined by the slower rate of the double-exponential function.

The probability of measuring a particular time of binding event is influenced by the bleaching and blinking properties of the fluorescent dye. To assess the influence of these processes, along with errors in detection, the bleaching-time distributions were measured independently under the same conditions using cells fixed with 4% (v/v) paraformaldehyde that blocks molecule movement. As before, a bleaching-time survival probability curve was fitted by a double-exponential function to extract exponential-time constants. The MukB-HaloTag bleaching time constant, *t*_*bleach*_, was measured to be 48.8 ± 8.3 s. The bleaching corrected binding-time was calculated by *t*_*bound*_ = *t*_*measured*_ * *t*_*bleach*_ / (*t*_*bleach*_ – *t*_*measured*_) (*34*). Blinking of fluorescent dye before or during binding events does not influence the measurement, because the observed binding times follow an exponential distribution and are therefore memoryless. All data analysis was performed in MATLAB (MathWorks).

#### 3D-structured illumination microscopy

Super-resolution 3D-structured illumination microscopy (SIM) images were acquired on a DeltaVision OMX V3 Blaze instrument (GE Healthcare), equipped with a 60×/1.42 oil UPlanSApo objective (Olympus), 405 nm, 488 nm and 593 nm diode lasers and three sCMOS cameras (PCO). Multiple-color three-dimensional stacks of MukB– Ypet/TetR-mCerulean were imaged sequentially. For each color, the raw 3D-SIM stacks were composed of 225 512×512 pixel images consisting of 21 z-sections (125 nm z-spacing, sample thickness of 2.5 mm). Each section consisted of 15 images - 3 angles and 5 phase shifts. Additionally, LacI-mCherry was imaged in a conventional wide-field mode. Acquisition settings were as follows: MukB–mYpet, 20 ms exposure with 488 nm laser (attenuated to 30% transmission); TetR-mCerulean, 50 ms exposure with 405 nm laser (30% transmission), LacI-mCherry, 50 ms exposure with 593 nm laser (30% transmission). The 3D-SIM raw data was computationally reconstructed with SoftWoRx 6.0 (GE Healthcare) using a Wiener filter setting of 0.004 and channel specific optical transfer functions to generate a super-resolution three-dimensional image stack with a lateral (x–y) resolution of ~120nm (wavelength dependent) and an axial (z) resolution of ~300 nm. In the reconstruction process, the pixel size was halved from 80 nm to 40 nm and the pixel number doubled in order to meet the Nyquist sampling criterion. For multichannel 3D alignment, mouse C127 cells were three-color (405, 488 and 594 nm) 5-ethenyl-2′-deoxyuridine (EdU) pulse-labeled as described in (*36*). The multichannel 3D-SIM EdU foci images were captured and reconstructed as described above, then channels were 3D corrected for chromatic shifts using the open-source software Chromagnon (github.com/macronucleus/ chromagnon)(*37*). The correction parameters obtained were then applied to align images from the experiments.

For analysis of MukBEF structure dimensions (Fig. 3D-F, Fig. S6), 3D-SIM image stacks were projected along the z-axis and the maximum intensity for each pixel selected using ImageJ. Only filaments with clear orientation in xy-axes were selected for analysis. Pixels belonging to the MukBEF structure were separated from the background using Otsu’s thresholding (*38*) in which the optimum threshold is chosen to minimize intra-class variance while maximizing inter-class variance. The structure’s centerline was calculated by using a morphological operation that erodes pixels from edges until only center pixels of the structure are left. Following this, branches of length 1 in the centerline were removed. The length of linear structures was measured as the minimum length of a curve that includes all pixels of the centerline. The length of circular filament was measured as the minimum contour length of a polygon that includes all pixels of the backbone. The thickness of the structure was measured by fitting a linegraph of pixel intensities crossing the centerline with a Gaussian function. Pixels close to the ends or branching points of the backbone were removed from the analysis. Further, the linegraph orientation was selected around a pixel to be normal to the structure so as to minimize width. From the Gaussian fit, full-width half-maximum (FWHM) distance was calculated as follows:

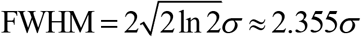

where σ is the standard deviation of the fitted Gaussian. All data analysis was performed in MATLAB (MathWorks).

### Simulations of loop extrusion

Stochastic simulations were performed using SGNS2 (*39*) (available https://sites.google.com/view/andreribeirolab/home/software), which uses the Gillespie method (Stochastic Simulation Algorithm) (*40*) to obtain exact realizations of the Chemical Master Equation (CME). SGNS2 supports dynamic compartments that can be created or destroyed during a simulation. The circular chromosome of *E. coli* was divided into 4641 discrete DNA segments, with each segment corresponding to a specific 1 kbp region of the chromosome. MukBEF is, unless otherwise stated, modelled as a dimer of dimers, which randomly binds to 2 adjacent free sites on the chromosome with a stochastic rate (k_bind_) with equal probability throughout the chromosome. Binding of MukBEF creates a dynamic compartment that contains a single DNA loop where each dimer of MukBEF occupies a single DNA segment. Following the binding event, each dimer of the compartment moves unidirectionally and independently away from each other one DNA segment at a time (releasing previous DNA segment while occupying the consecutive one) with a stochastic rate (k_move_) for extrusion of a loop. As the MukBEF dimers move away, DNA in the loop is free to be bound by other MukBEF molecules allowing for the creation of loops inside loops. If dimers collide on the chromosome head-on, they block each other, while the other dimers of the compartment continue loop extrusion unperturbed. Unbinding of MukBEF releases the DNA segments under its footprint and destroys the loop with a stochastic rate (k_unbind_) that is independent of the state of the chromosome or other MukBEF. The residency time is the same everywhere on the chromosome, except in simulations with displacement of MukBEF from *ter* region, where binding or moving leads to instant dissociation of MukBEF molecule and destruction of the loop. Aforementioned reactions of the model are written for every DNA segment of the system except in ter region where an additional reaction releases MukBEF upon contact. Asymmetric loop extrusion is modelled by only one of the dimers moving away from the binding site orientation decided randomly at the binding. The cytosolic state of MukBEF is assumed well-mixed and is therefore treated implicitly.

The rate constants were used as measured here. Namely, the MukBEF unbinding rate (k_unbind_) is 0.0154 s^−1^ and the MukBEF binding rate (k_bind_) is 3.9e-06 s^−1^ per DNA segment per free MukBEF complex and was adjusted to result in 48% of MukBEF to be bound to the chromosome with wild type MukBEF copy numbers (110 MukBEF dimer of dimers)(*2*). The loop extrusion rate (k_move_) has not been measured *in vivo* and therefore was set to 0.6 DNA segments/s/dimer (corresponding to 600 bp/dimer/s) as estimated *in vitro* (*20*). The expected loop size without collisions is 80 kbp (40 kbp for unidirectional loop extrusion). The system state including the state of each loop compartment was read out after 500 s to allow the overall loop structure on the chromosome to reach maturation. Each simulation was repeated at least 1000 times to ensure proper sampling of chromosome states.

In the analysis of simulated chromosomes, MukBEF clusters were defined as MukBEF molecules that do not have empty DNA segments between them. Loops inside loops can contribute to the cluster size if they have reached the stem of the main loop by at least one dimer. After finding the largest MukBEF cluster, the shortest distance between the largest cluster and a chromosome locus was measured along the chromosome from a DNA segment of the specific chromosome locus to the closest MukBEF of the largest cluster. Loops acts as ‘shortcuts’ decreasing the distances between chromosomal loci. The loop state of a single chromosome (Fig. 4C) was shown as a polar coordinate plot that shows the starting and the ending locations of DNA loops or as a 2D force-directed layout of the circular chromosome after converting the loop state into a graph with loops as connections between otherwise circular organization of DNA segments. Random links (Fig. 4A) were generated by adding 110 (wild type occupancy) connections between random DNA segments. All data analysis was performed in MATLAB (MathWorks).

---

**Fig. S1.**
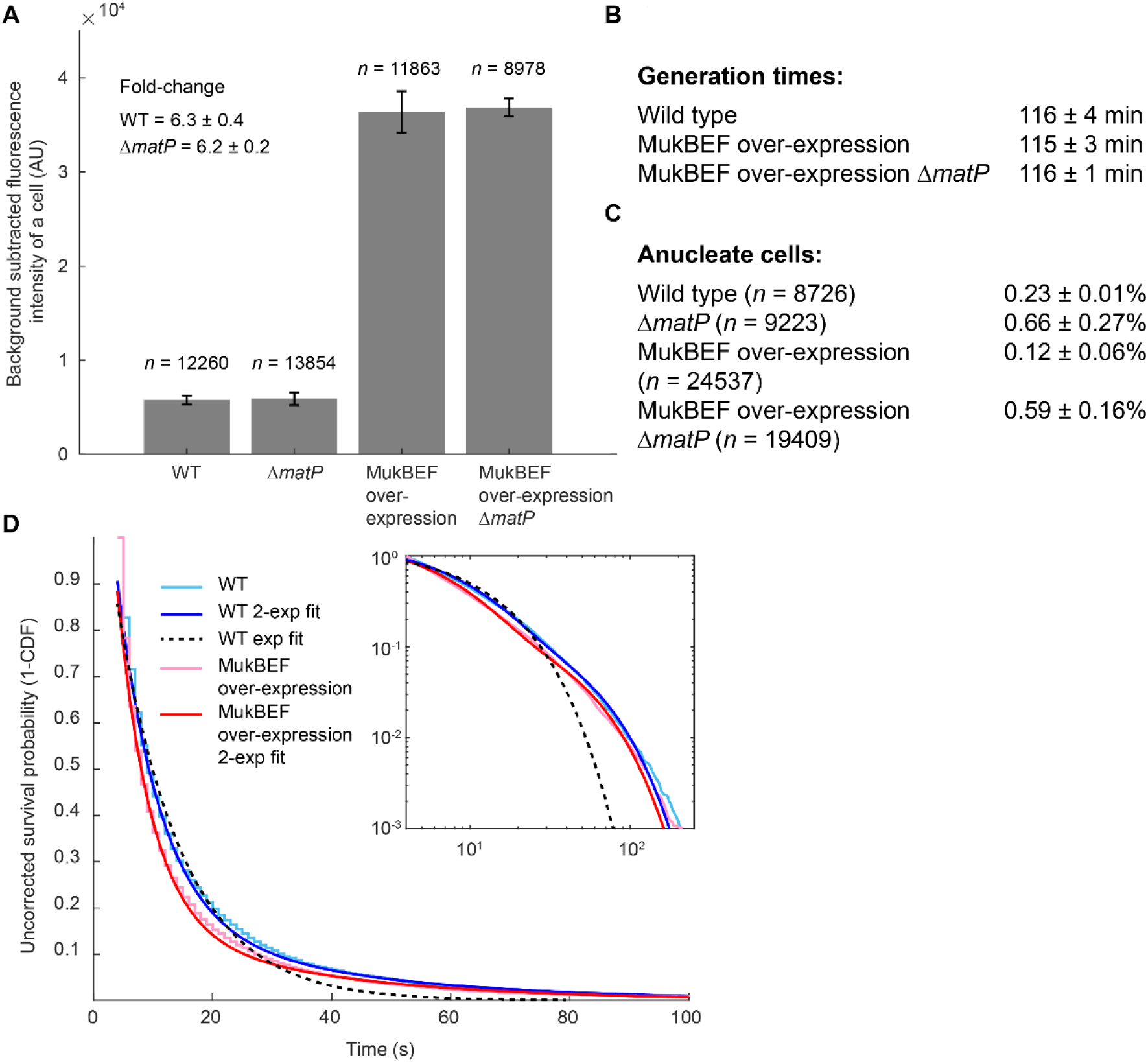
Characterization of cells with increased MukBEF chromosome occupancy. (**A**) MukB-mYpet fluorescence intensity from 3 repeats in WT, Δ*matP*, MukBEF over-expression and MukBEF over-expression Δ*matP* cells and fold-change between WT and MukBEF over-expression. (**B**) Generation times from 3 repeats (±SEM) for WT, over-expression and over-expression Δ*matP*. (**C**) Anucleate cell percentages from 3 repeats (±SEM) in WT, Δ*matP*, MukBEF over-expression and MukBEF over-expression Δ*matP*. (**D**) Uncorrected survival probability (1-CDF) curves of track lengths measured from single molecule tracking of MukB^JF549^ with a 1 s exposure time in wild type (10084 tracks; 4 experiments) and increased occupancy (14734 tracks; 4 experiments). The experimental data are fitted by a double-exponential functions. For comparison, an exponential fit to wild type is shown. Inset shows log-log plot of the same data. The bleaching rate was measured to be 48.8 ± 8.3 s (9739 tracks; 3 experiments). The bleaching corrected residency time for wild type cells is 66.9 ± 15.3 s and for increased occupancy cells 63.7 ± 13.5 s.

**Fig. S2.**
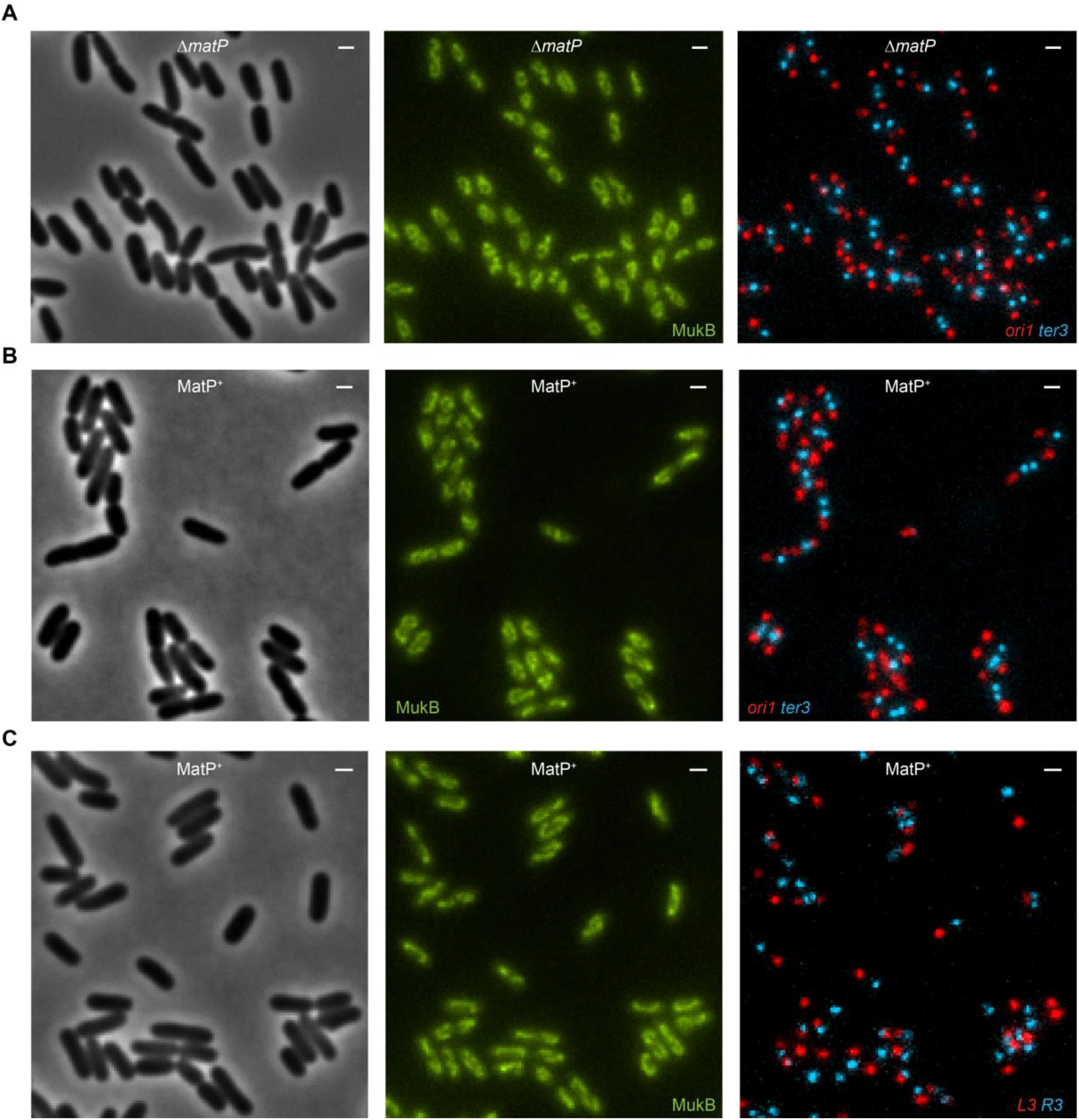
MukBEF increased occupancy cells in MatP^+^ (with MatP present) and ΔmatP. Representative phase contrast and fluorescence images of cells with (**A**) MukBEF increased occupancy Δ*matP* cells with *ori1* and *ter3* markers, (**B**) MukBEF increased occupancy with *ori1* and *ter3* markers, and (**C**) MukBEF increased occupancy with *L3* and *R3* markers. Scale bars, 1 μm.

**Fig. S3.**
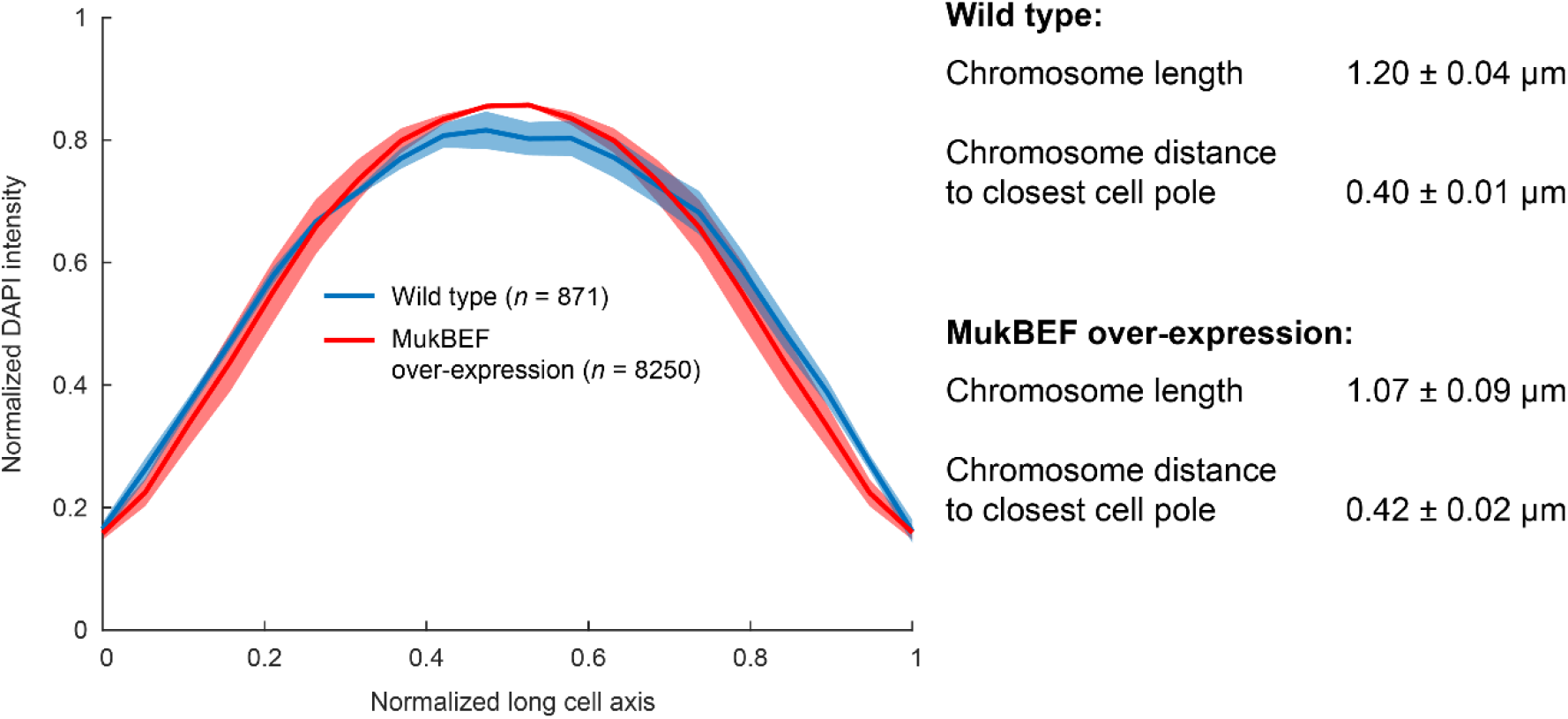
DAPI profiles in WT and MukBEF increased occupancy cells. Normalized DAPI intensity profiles on normalized long cell axis for WT (871 cells) and MukBEF increased occupancy (8250) cells. Only cells below 2.6 μm long were considered to avoid cells with more than 1 chromosomes. DAPI length was measured as full-width-half-maximum (FWHM) of the DAPI profile. Also, the distance to the cell pole from half maximum of the DAPI profile was measured. Data are from 3 repeats (±SEM).

**Fig. S4.**
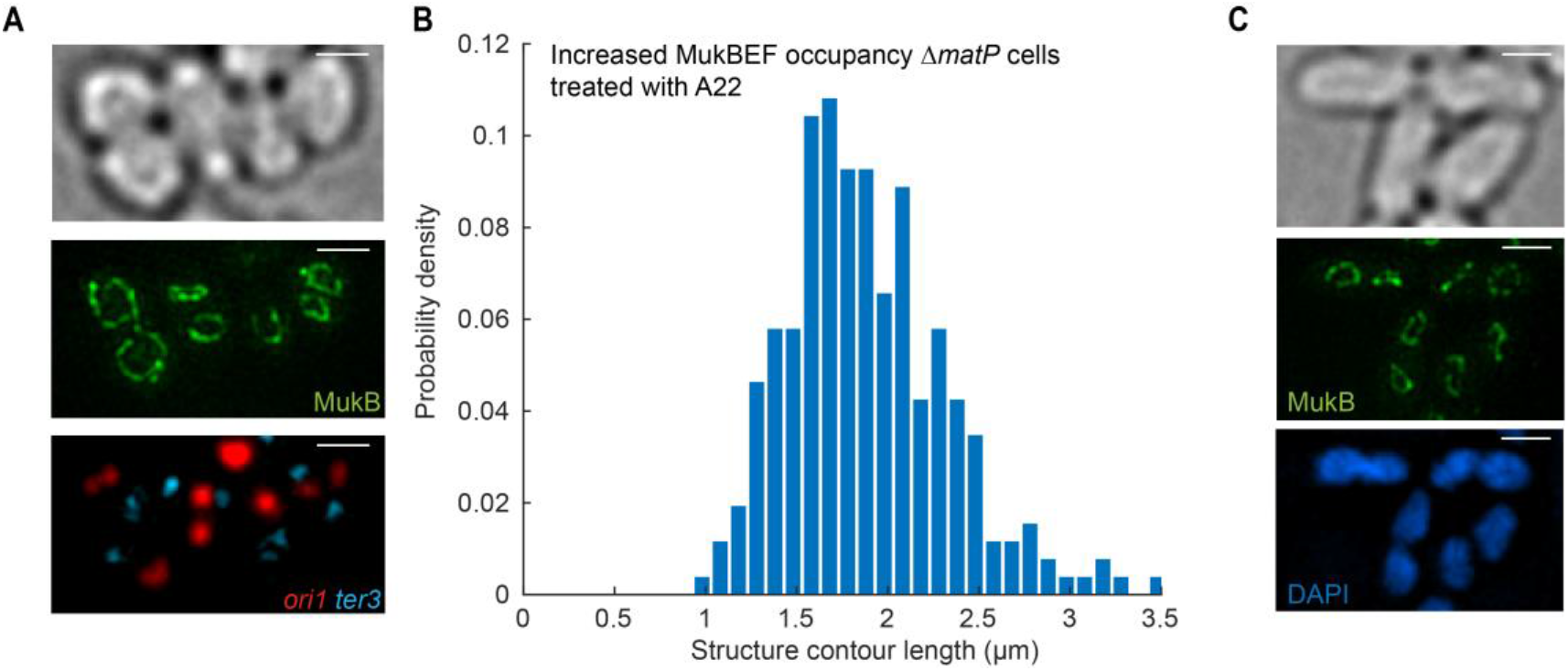
MukBEF increased occupancy cells treated with A22 or rifampicin. (**A**) Representative SIM images of MukBEF increased occupancy Δ*matP* cells with *ori1* and *ter3* markers grown with A22 (4 μg/ml) for 6 h. 3D-SIM images were maximum projected onto 2D for visualization purposes. Scale bars, 1 μm. (**B**) Distribution of axial core contour lengths (*n* = 259) measured from the cells in (A). (**C**) Representative SIM images of MukBEF increased occupancy Δ*matP* cells with *ori1* and *ter3* markers grown with rifampicin (0.025 μg/ml) for 2 h. Scale bars, 1 μm.

**Fig. S5.**
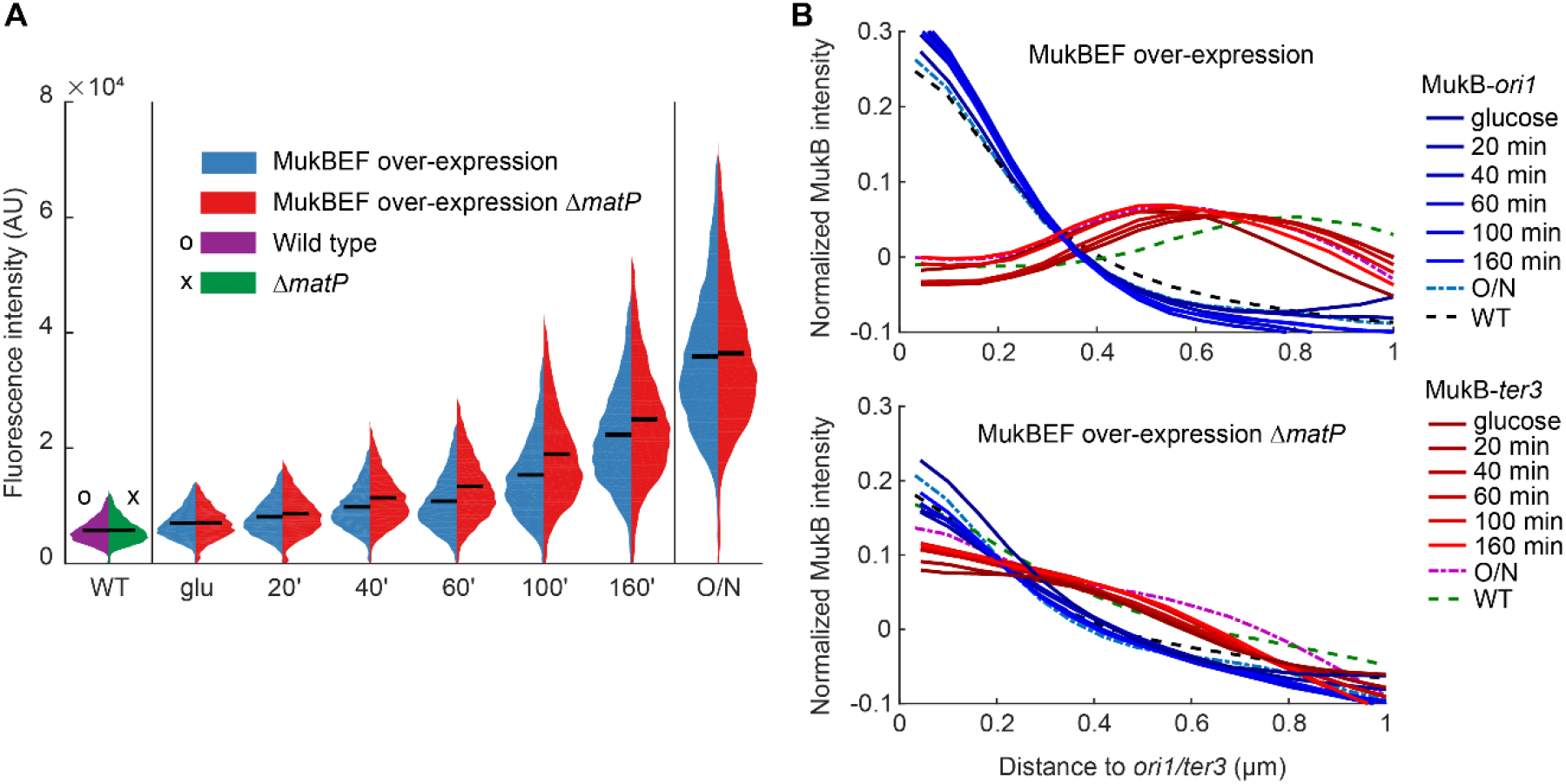
Induction of MukBEF over-expression time series. (**A**) MukB-mYpet intensity in wild type and MukBEF over-expression strain induction with 0.2 % (w/v) arabinose. Fluorescence expression levels are shown prior to induction (glucose), during induction (20 min, 40 min, 60 min, 100 min, 160 min) and cultures grown over night in the presence of arabinose. Left distributions correspond to MukBEF increased occupancy MatP^+^ strain and right distributions to the MukBEF increased occupancy Δ*matP* strain. Also shown are wild type and Δ*matP* strains. Black lines show the means. Data are from at least 2 independent experiment with number of data points in WT (12260, 13854), glucose (9537, 4614), 20 min (8521, 9385), 40 min (11419, 5939), 60 min (8982, 8995), 100 min (9630, 10152), 160 min (9396, 10212), and over-night induction (11863, 8979) in MatP^+^ and Δ*matP* cells, respectively. (**B**) Normalized MukB-mYpet pixel intensity as a function of distance to *ori1*/*ter3* in (top) MukBEF increased occupancy and (bottom) MukBEF increased occupancy Δ*matP* cells. Also shown are wild type and Δ*matP* strain. Data are same as in (A).

**Fig. S6.**
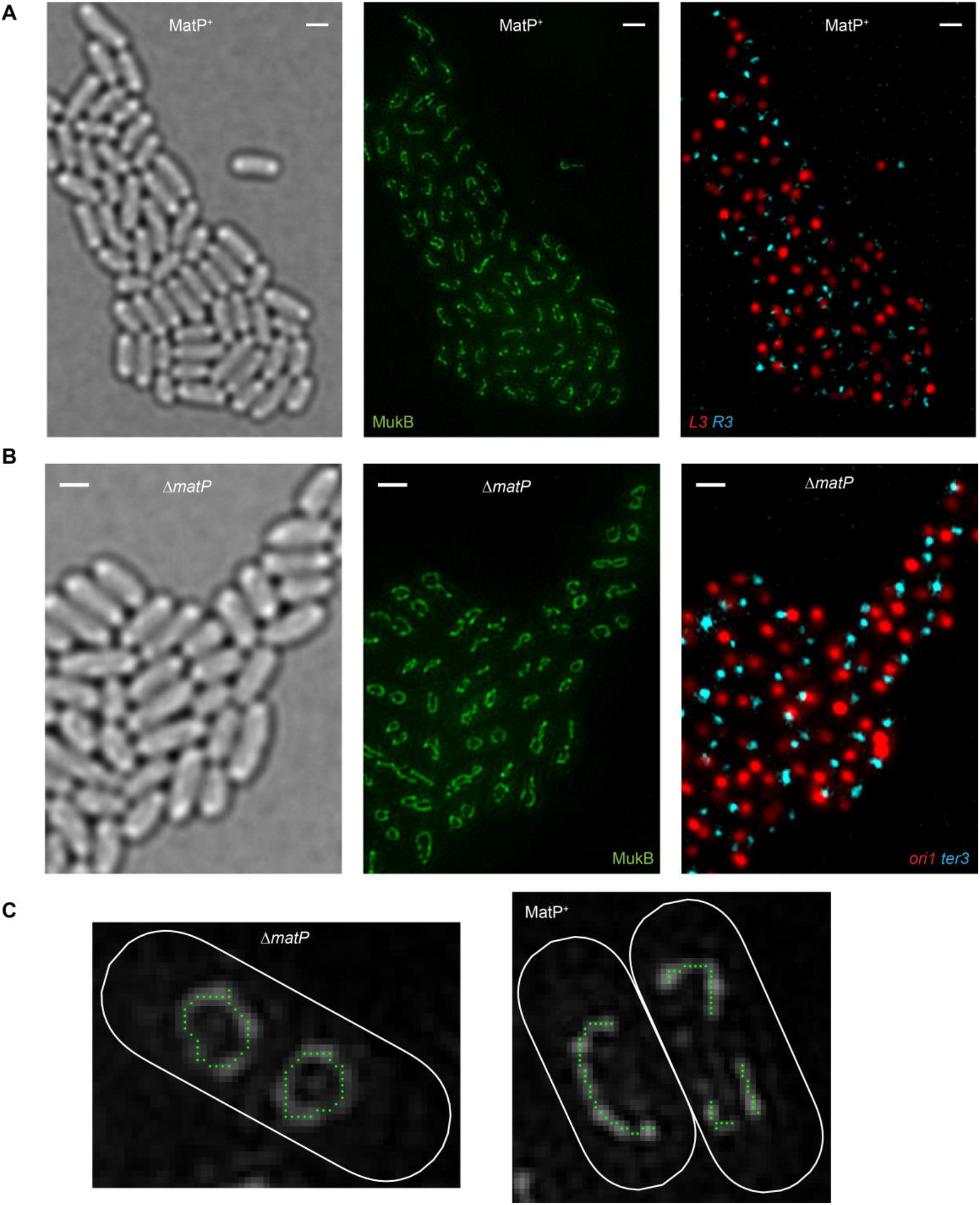
SIM images and MukBEF structure length analysis. Representative SIM images of MukBEF increased occupancy in (**A**) MatP^+^ (with MatP present) cells with *L3* and *R3* markers, and (**B**) Δ*matP* cells with *ori1* and *ter3* markers. 3D-SIM images were maximum projected onto 2D for visualization. Scale bars, 1 μm. (**C**) Example images of detected centerlines of circular and linear the MukBEF structures in Δ*matP* and MatP^+^ cells, respectively. White line is the cell border. Green dots are centerline pixels of the structure.

**Fig. S7.**
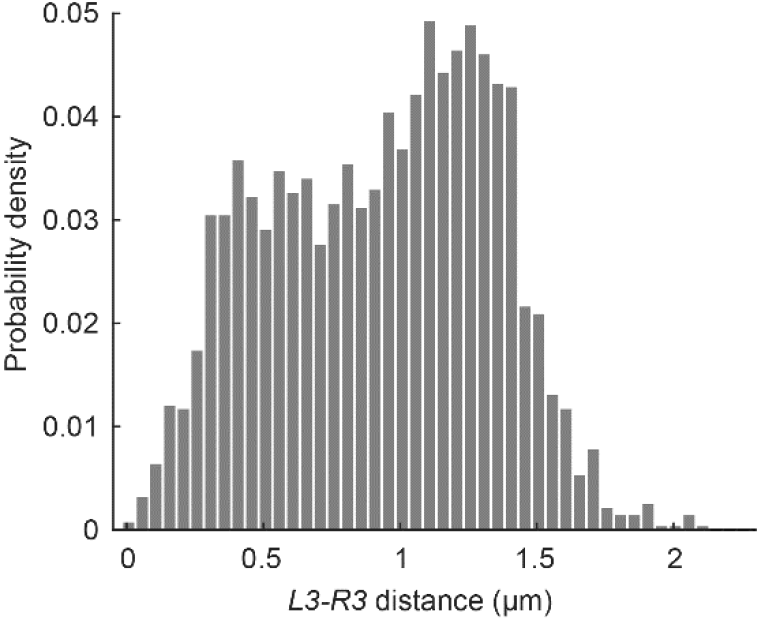
*L3-R3* distances in MukBEF increased occupancy cells. Distances between *L3* and *R3* markers in MukBEF increased occupancy cells. 2824 single chromosome cells from 2 experiments were analyzed.

**Fig. S8.**
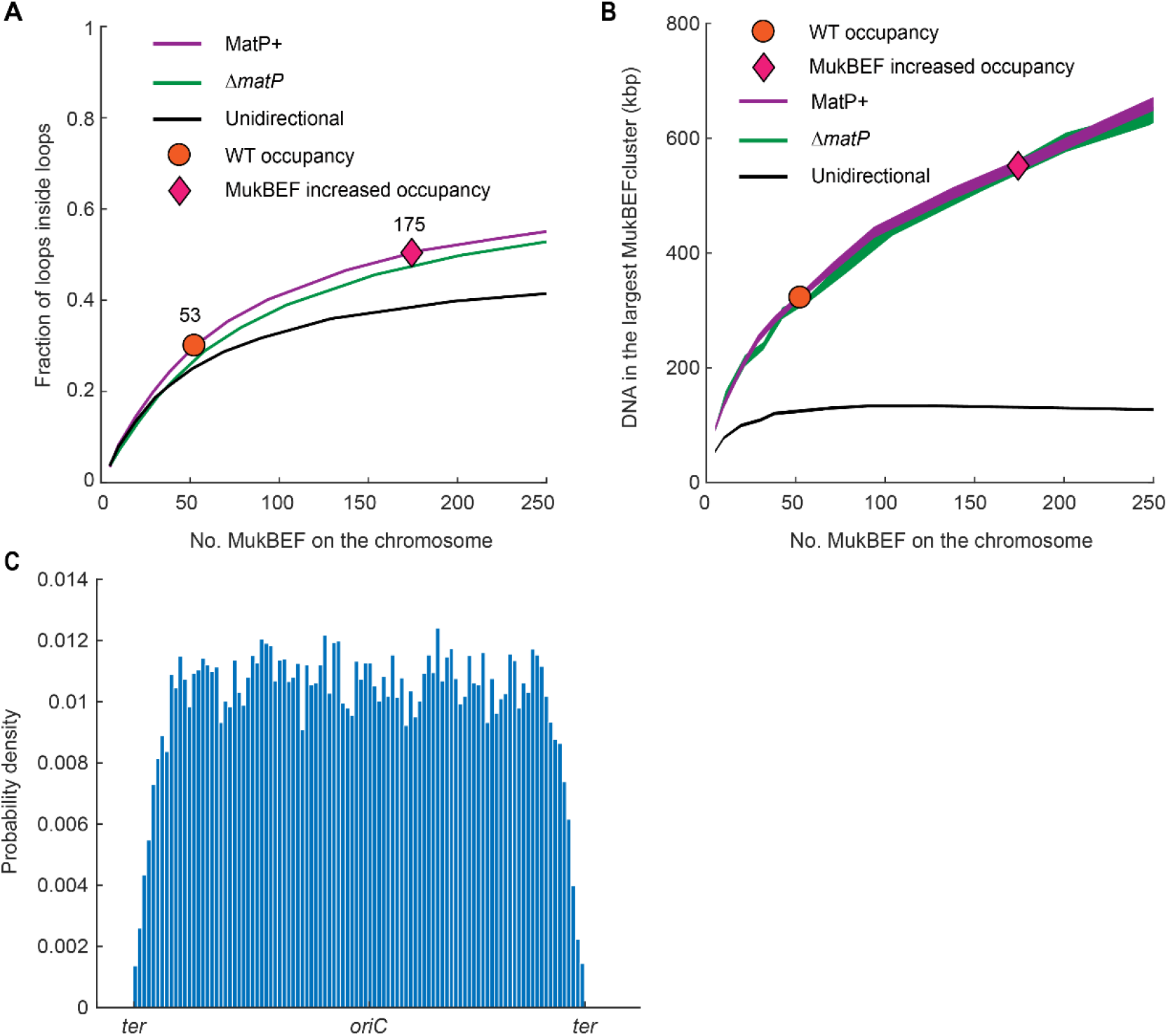
Modeling loop extrusion by MukBEF. (**A**) Fraction of loops inside loops and (**B**) DNA in the largest MukBEF cluster (no gaps between MukBEF) as function of the number of bound MukBEF dimers of dimers. Experimentally observed numbers of MukBEF dimers of dimers on the chromosome for wild type (○) and MukBEF increased occupancy (◊) are depicted. Additionally, unidirectional model of loop extrusion (black line) is shown where each dimers binds and extrudes a loop independently in a randomly chosen direction. Line thickness denotes 95% bootstrap confidence interval for the mean across at least 1000 simulation replicas. (**C**) MukB occupancy profile on the chromosome with wild type MukBEF occupancy across 4000 simulation replicas.

**Table S1.** Strain list. *kan*, *cat*, *gen*, and *hyg* refer to insertions conferring resistance to kanamycin (Km^r^), chloramphenicol (Cm^r^), gentamycin (Gm^r^) and hygromycin B (Hyg^r^), respectively. *frt* refers to the Flp site-specific recombination site.

| Strain | Relevant genotype | Source or reference |
| --- | --- | --- |
| AB1157 | F<sup>-</sup>, λ<sup>-</sup>, *rac*<sup>-</sup>, *thi-1*, *hisG4*, Δ(*gpt-proA*)62, *argE3*, *thr-1*, *leuB6*, *kdgK51*, *rfbD1*, *araC14*, *lacY1*, *galK2*, *xylA5*, *mtl-1*, *tsx-33*, *supE44(glnV44)*, *rpsL31(str<sup>R</sup>)*, *qsr'-0*, *mgl-51* | Coli Genetic Stock Center (CGSC) #1157 (27) |
| SN192 | AB1157, *lacO240* at *ori1* (3908) (*hyg*), *tetO240* at *ter3* (1644) (*gen*), Δ*leuB::Plac-lacI-mCherry-frt*, Δ*galk::Plac-tetR-mCerulean-frt*, *mukB-Ypet-frt* | (8) |
| SN302 | AB1157, *lacO240* at *ori1* (3908) (*hyg*), *tetO240* at *ter3* (1644) (*gen*), Δ*leuB::Plac-lacI-mCherry-frt*, Δ*galk::Plac-tetR-mCerulean-frt*, *mukB-Ypet-frt*, Δ*matP::cat* | (8) |
| JM41 | AB1157, *mukB-HaloTag-frt-kan-frt* | (15) |
| JM56 | AB1157, *mukB-HaloTag-frt*, Δ*mukE::frt-kan-frt* | This work |
| JM90 | AB1157, *lacO240* at *ori1* (3908) (*hyg*), *tetO240* at *ter3* (1644) (*gen*), Δ*leuB::Plac-lacI-mCherry-frt*, Δ*galk::Plac-tetR-mCerulean-frt*, *frt-kan-frt-araC-Para-smtA-mukFE-mukB-Ypet-frt* | This work |
| JM91 | AB1157, *lacO240* at *ori1* (3908) (*hyg*), *tetO240* at *ter3* (1644) (*gen*), Δ*leuB::Plac-lacI-mCherry-frt*, Δ*galk::Plac-tetR-mCerulean-frt*, *frt-kan-frt-araC-Para-smtA-mukFE-mukB-Ypet-frt*, Δ*matP::cat* | This work |
| JM101 | AB1157, *lacO240* at *L3* (2268) (*hyg*), *tetO240* at *R3* (852) (*gen*), Δ*leuB::Plac-lacI-mCherry-frt*, Δ*galk::Plac-tetR-mCerulean-frt*, *frt-kan-frt-araC-Para-smtA-mukFE-mukB-Ypet-frt* | This work |
| JM103 | AB1157, *frt-araC-Para-smtA-mukFE-mukB-HaloTag-frt-kan-frt* | This work |

**Table S2.**
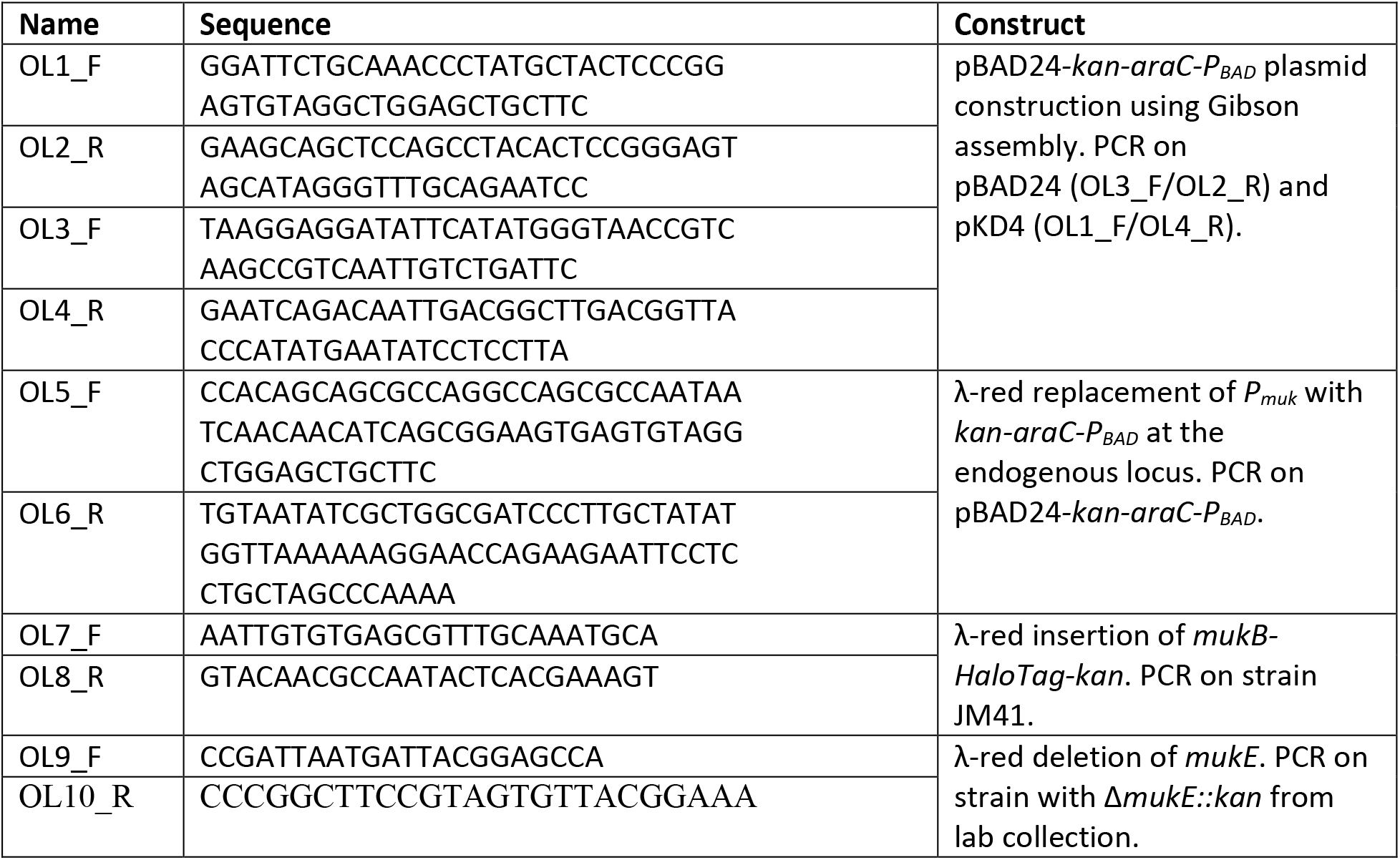
Primer list.

